# Comparison of black soldier fly, cricket, and superworm on growth performance, nutrient utilization and fatty acid profiles of rainbow trout

**DOI:** 10.1101/2024.07.15.603309

**Authors:** Sonja Drosdowech, Marcia Chiasson, David Ma, David Huyben, Neil Rooney

## Abstract

Inclusion of fishmeal and fish oil in feeds for farmed fish is not sustainable and alternatives need to be evaluated. The aim of this study was to determine the effect of diets including defatted black soldier fly larvae (*Hermetia illucens*), adult cricket (*Gryllodes sigillatus*) and superworm (*Zophobas morio*) on the growth performance, apparent nutrient digestibility, nutrient retention, and fatty acid profiles of juvenile rainbow trout (*Oncorhynchus mykiss*). Triplicate tanks of fish (100.5 ± 0.6 g; mean ± SD) were fed one of four diets, a control diet with 20% fishmeal, and three experimental diets containing either 15% defatted black soldier fly meal, 15% full-fat adult cricket meal or 15% full-fat superworm meal, where each insect meal partially replaced fishmeal and fish oil. After 84 days of feeding, no significant differences were observed between diets for growth performance indicators or body indices. Fish fed the control and superworm diets had a higher protein content and retention in whole-body carcass compared to the cricket and black soldier fly diet groups. No significant effects between diets were found on whole-body carcass regarding fatty acid classes SFA, MUFA, PUFA n-3 and PUFA n-6, although lauric acid and myristic acid were significantly higher in fish fed black soldier fly diet and linoleic acid was higher in fish fed superworm and cricket diets. Retention of fatty acids was higher for most classes in fish fed the black soldier fly diet, yet whole carcass lipid content did not differ significantly between diets. Additionally, apparent digestibility of phosphorous was significantly improved for all insect diets compared to the control. These results indicate that insect meals can partially replace fishmeal and fish oil in diets for rainbow trout without compromising growth performance and fatty acid composition, while improving phosphorous utilization.

## 1. Introduction

The escalating demand for protein, driven by a growing human population and changing dietary preferences, has placed immense pressure on global food production systems. Fish are a vital source of protein and omega-3 fatty acids for humans and demand is growing due to their numerous health benefits and nutritional value (FAO, 2022). Since the plateau of fish production from wild fisheries during the 1990’s, increased demand for fish protein has been met by an expanding aquaculture industry (FAO, 2022). However, carnivorous fishes that require high protein and high oil diets, such as Atlantic salmon (*Salmo salar*) and rainbow trout (*Oncorhynchus mykiss*), have a heavy reliance on wild-caught fish protein and oil in the diet (Tacon & Metian, 2015). While the total content of fishmeal and oil used in aquafeeds has decreased over time, the aquaculture industry has continued to grow leading to continued high demand of these ingredients (Jannathulla et al., 2019; Naylor et al., 2021). Landings of forage fish that are used to make fish meal and fish oil have decreased in recent years, while the price has more than doubled during the 2000s (Naylor et al., 2021). In response to the increasing price, and reduced ability of wild capture fisheries to meet this demand, the aquaculture industry has looked towards nutritionally viable feed ingredients.

The substitution of fishmeal and fish oil with insect meal has emerged as a promising solution to address the sustainability challenges associated with protein and lipid sourcing in aquaculture. Insects are efficient at converting organic waste into a valuable food source, high in protein and lipids (Sánchez-Muros et al., 2014). Insect meal is also hypothesized to offer potential health benefits to fish that result in improved growth performance (Barroso et al., 2014; Hossain et al., 2021; Rema et al., 2019). Additionally, insect meal has been proposed as an environmentally sustainable substitute for fishmeal and fish oil, providing an opportunity to mitigate ecological concerns associated with the reliance on wild-caught fish for food animal production (Alfiko et al., 2022). Commercialization of insect ingredients in aquaculture requires further research addressing the effects of insect feed on the health and productivity of cultured fish, as well as the potential impacts on product quality.

Most research focused on replacing fishmeal with insect meal in the diets of rainbow trout has focused on black soldier fly (*Hermetia illucens*) and yellow mealworm (*Tenebrio molitor*) (Belforti et al., 2015; Józefiak, Nogales-Mérida, Mikołajczak, et al., 2019; Rema et al., 2019; Rimoldi et al., 2021). In addition to improving the gut microbiota of rainbow trout, recent studies have found that black soldier fly may be included in the diet at up to 30% without compromising fish growth performance (Huyben et al., 2019; Rimoldi et al., 2019; Terova et al., 2019). Yellow mealworm has been included at up to 50% in the diet of rainbow trout without compromising growth and has been observed to improve feed conversion (Belforti et al., 2015). Further, dietary inclusion of 25% yellow mealworm has been shown to improve crude protein, phosphorous and gross energy retention in rainbow trout (Rema et al., 2019). However, black soldier fly and mealworm contain a high content of lipid and are usually defatted to increase crude protein content, which is still moderate around 50-55% compared to 60-75% for fishmeal (Gasco et al., 2019). Therefore, other insect meals need to be investigated as alterative ingredients in rainbow trout diets.

Currently, alternative insect species such as adult cricket (*Gryllodes sigillatus*), and superworm (*Zophobas morio*) have not been widely examined as a dietary supplement for rainbow trout. One study on the use of superworm meal in rainbow trout production found that while 11% inclusion in the diet did not significantly affect growth performance, 22% inclusion had negative impacts on fish growth and feed conversion (Shekarabi et al., 2021). With respect to crickets, Józefiak et al. (2019) found that a 20% inclusion rate of full fat cricket meal negatively impacted rainbow trout growth performance. Interestingly, it has been observed that live superworm and crickets, and a combination of the two insects can feasibly replace a commercial diet of the same energy content without negatively impacting rainbow trout growth performance (Turek et al., 2020). These few studies, while valuable, leave questions regarding the relative effects that different insect meals may have on rainbow trout growth performance when compared to fish meal diets.

Studies examining the inclusion of insect meal in diets for rainbow trout have documented significant effects on the whole-body composition, including alterations in the fatty acid profile (Bruni et al., 2020; Fabrikov et al., 2021; Mancini et al., 2018; Turek et al., 2020). The fatty acid composition of insect meal varies depending on the species and rearing substrate (Chieco et al., 2019; Danieli et al., 2019; Georgescu et al., 2022; Meneguz et al., 2018; Riekkinen et al., 2022), leading to potential implications on the health of rainbow trout and humans that consume these fish. For example, black soldier fly possesses relatively higher levels of saturated fatty acids and lower amounts of essential n-3 long-chain polyunsaturated fatty acids (LC-PUFA) compared to fish meal (Sánchez-Muros et al., 2014). These characteristics could be perceived as less desirable for consumers who seek fish products primarily for their high content of health-promoting n-3 LC-PUFA. Recent studies have also found that feeding insect meal to rainbow trout results in fatty acid profile characterized by decreased levels of n-3 LC-PUFAs in their fillets (Fabrikov et al., 2021; Mancini et al., 2018; Turek et al., 2020). However, most studies to this point have focused on the effects of varying dietary inclusion levels of black soldier fly on rainbow trout fatty acid profiles, overlooking the potential for alternate species to serve as viable fish meal substitutes. Comparing the specific changes that different insect meals induce in rainbow trout fatty acid composition is crucial for ensuring the production of high-quality, nutritionally balanced fillets that meet the expectations and preferences of health-conscious consumers.

The aim of this study was to determine the effects of partially replacing fishmeal and fish oil with black soldier fly, adult cricket, and superworm meals on growth performance, nutrient retention, apparent nutrient digestibility, and fatty acid composition of juvenile rainbow trout. Insect meal diets were formulated to be as similar in protein, fat and energy as possible while comprising 15% of the total diet.

## 2. Materials and Methods

### 2.1 Experimental Fish and Husbandry Conditions

Fish were maintained in accordance with animal utilization protocol #4794, reviewed and approved by the University of Guelph’s Animal Care Committee. Juvenile mixed sex rainbow trout (N= 600) at 100.5 ± 0.6 g (mean ± SD) were hatched and raised at the Ontario Aquaculture Research Centre (OARC; Elora, ON, Canada). Fish were randomly distributed in groups of 50 to 12 fiberglass tanks (330 L) supplied with aerated and degassed groundwater at 20-25 L/min. Before initiating the feeding trial, fish were acclimated for 14 days in the experimental tanks and were fed a standard commercial diet (Bluewater Trout 3mm, Sharpe Farm Supply, Guelph, ON, Canada). Water quality was measured weekly for temperature (8.5 ± 0.1°C) and dissolved oxygen (≥ 70 % saturation). Fish were maintained under a simulated natural photoperiod for 43.5° North using LED lights and photoperiod controllers to simulate dawn and dusk.

Each of the four experimental diets were randomly assigned to three replicates tanks of the 50 juvenile rainbow trout using a randomized block design. Individual fish weight and fork length of all fish were recorded to determine initial growth parameters. During the 12-week trial, fish were hand fed to apparent satiation twice per week in two separate feed offerings (9-12:00 and 13-16:00 h). On the remaining 5 days of the week, 95% restriction of the two prior hand feeding events was fed using automatic belt feeders. Feed intake was recorded daily.

### 2.2 Experimental Diet Formulation

Superworm meal *(Zophobas morio)* and cricket meal *(Gryllodes sigillatus)* were provided by Entomo Farms (Norwood, ON, Canada), while black soldier fly meal *(Hermetia illucens)* was provided by Enterra Feed Corporation (Langley, BC, Canada). Samples of fishmeal, and the three insect meals were analyzed for proximate and fatty acid composition (Table 1).

**Table 1.**
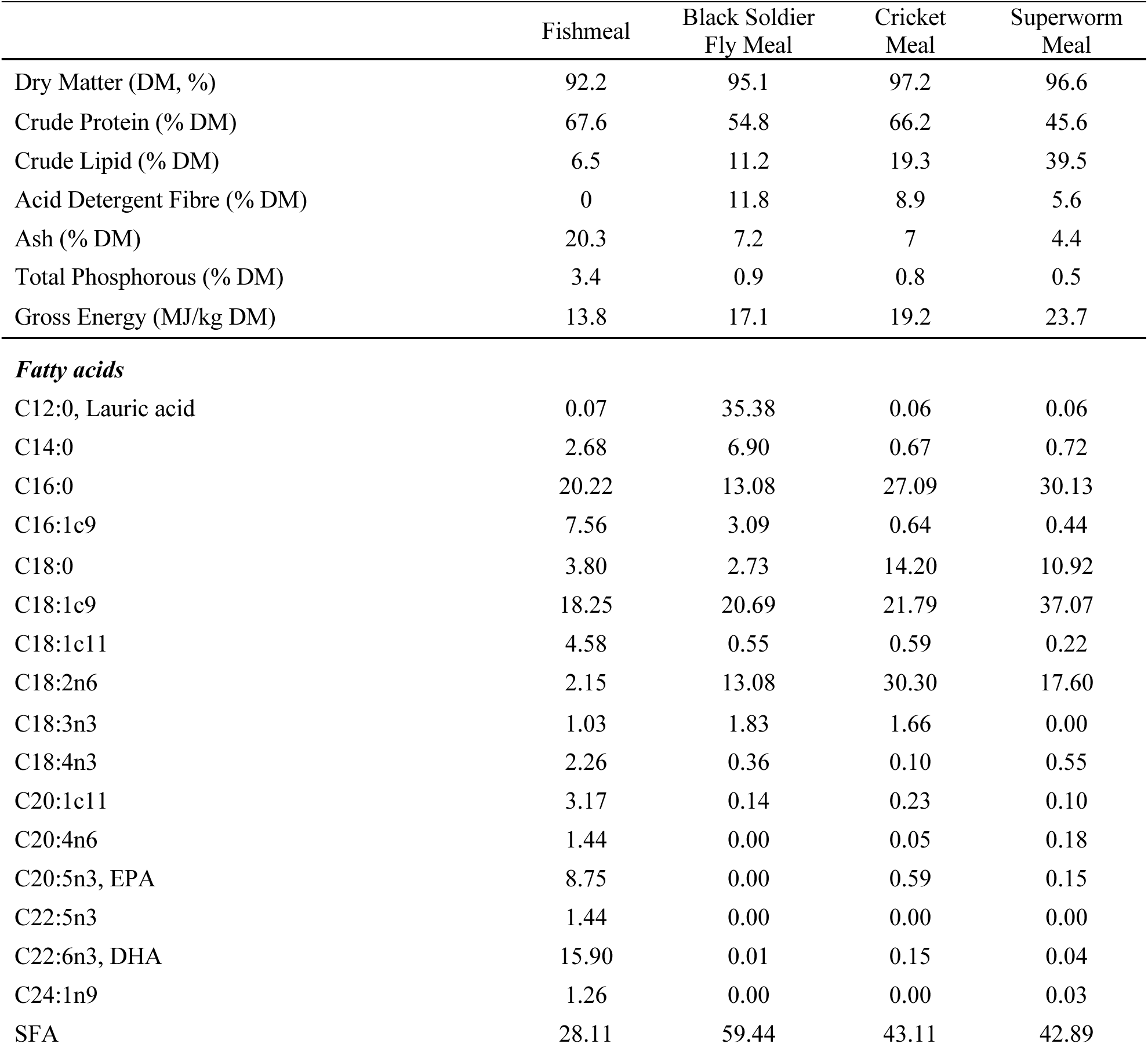
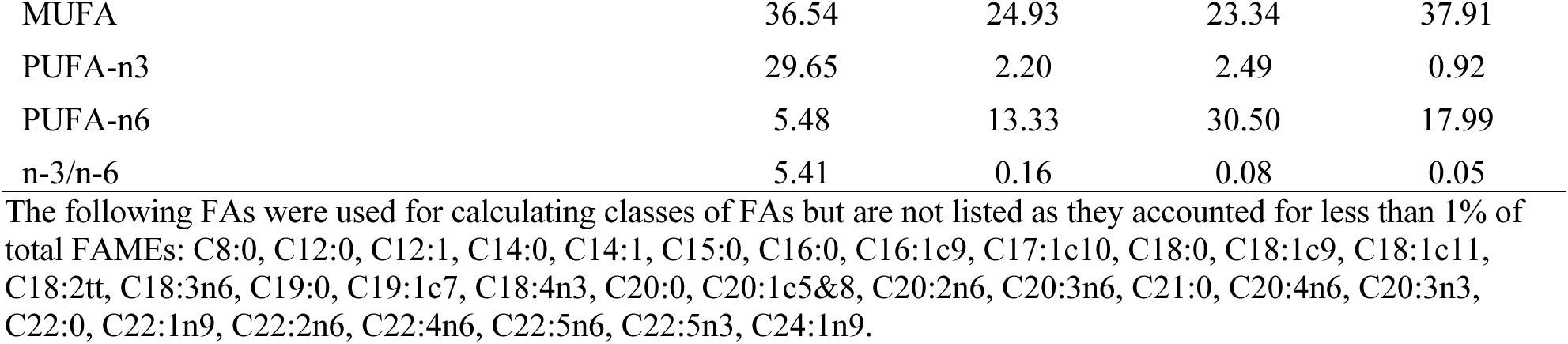
Proximate composition and fatty acid profile reported as % of total Fatty Acid Methyl Esters (FAMEs) of fishmeal and insect meal ingredients.

|  | Fishmeal | Black Soldier Fly Meal | Cricket Meal | Superworm Meal |
| --- | --- | --- | --- | --- |
| Dry Matter (DM, %) | 92.2 | 95.1 | 97.2 | 96.6 |
| Crude Protein (% DM) | 67.6 | 54.8 | 66.2 | 45.6 |
| Crude Lipid (% DM) | 6.5 | 11.2 | 19.3 | 39.5 |
| Acid Detergent Fibre (% DM) | 0 | 11.8 | 8.9 | 5.6 |
| Ash (% DM) | 20.3 | 7.2 | 7 | 4.4 |
| Total Phosphorous (% DM) | 3.4 | 0.9 | 0.8 | 0.5 |
| Gross Energy (MJ/kg DM) | 13.8 | 17.1 | 19.2 | 23.7 |
| ***Fatty acids*** |  |  |  |  |
| C12:0, Lauric acid | 0.07 | 35.38 | 0.06 | 0.06 |
| C14:0 | 2.68 | 6.90 | 0.67 | 0.72 |
| C16:0 | 20.22 | 13.08 | 27.09 | 30.13 |
| C16:1c9 | 7.56 | 3.09 | 0.64 | 0.44 |
| C18:0 | 3.80 | 2.73 | 14.20 | 10.92 |
| C18:1c9 | 18.25 | 20.69 | 21.79 | 37.07 |
| C18:1c11 | 4.58 | 0.55 | 0.59 | 0.22 |
| C18:2n6 | 2.15 | 13.08 | 30.30 | 17.60 |
| C18:3n3 | 1.03 | 1.83 | 1.66 | 0.00 |
| C18:4n3 | 2.26 | 0.36 | 0.10 | 0.55 |
| C20:1c11 | 3.17 | 0.14 | 0.23 | 0.10 |
| C20:4n6 | 1.44 | 0.00 | 0.05 | 0.18 |
| C20:5n3, EPA | 8.75 | 0.00 | 0.59 | 0.15 |
| C22:5n3 | 1.44 | 0.00 | 0.00 | 0.00 |
| C22:6n3, DHA | 15.90 | 0.01 | 0.15 | 0.04 |
| C24:1n9 | 1.26 | 0.00 | 0.00 | 0.03 |
| SFA | 28.11 | 59.44 | 43.11 | 42.89 |
| MUFA | 36.54 | 24.93 | 23.34 | 37.91 |
| PUFA-n3 | 29.65 | 2.20 | 2.49 | 0.92 |
| PUFA-n6 | 5.48 | 13.33 | 30.50 | 17.99 |
| n-3/n-6 | 5.41 | 0.16 | 0.08 | 0.05 |

The following FAs were used for calculating classes of FAs but are not listed as they accounted for less than 1% of total FAMES: C8:0, C12:0, C12:1, C14:0, C14:1, C15:0, C16:0, C16:1c9, C17:1c10, C18:0, C18:1c9, C18:1c11, C18:2tt, C18:3n6, C19:0, C19:1c7, C18:4n3, C20:0, C20:1c5&8, C20:2n6, C20:3n6, C21:0, C20:4n6, C20:3n3, C22:0, C22:1n9, C22:2n6, C22:4n6, C22:5n6, C22:5n3, C24:1n9.

The four experimental diets (three insect meal and one fishmeal control) were formulated to be isonitrogenous (46% crude protein), isolipidic (18% crude lipid), and isoenergetic (22 MJ/kg), meeting all known nutritional requirements for juvenile rainbow trout (NRC, 2011) (Table 2). Each insect diet formulation had an insect meal inclusion at 15% where fishmeal was replaced on a crude protein basis and fish oil as well as canola oil were replaced on a crude lipid basis. Corn starch inclusion was adjusted to achieve equal gross energy levels across all four diets. Additionally, 150 mg/kg yttrium oxide was added to each diet as an inert marker to calculate apparent digestibility of diets and feed ingredients.

**Table 2.** Formulation of the experimental diets.

| **Ingredient (%)** | **Control** | **Black Soldier Fly Meal** | **Cricket Meal** | **Superworm Meal** |
| --- | --- | --- | --- | --- |
| Fish meal, herring | 20 | 8.39 | 5.32 | 10.7 |
| Poultry by-product meal | 15 | 15 | 15 | 15 |
| Wheat flour | 15 | 15 | 15 | 15 |
| Soy protein, HP300 | 11 | 11 | 11 | 11 |
| Soybean meal | 5 | 5 | 5 | 5 |
| Black soldier fly larvae, defatted, Enterra |  | 15 |  |  |
| Cricket meal, Entomo |  |  | 15 |  |
| Superworm meal, full-fat, Entomo |  |  |  | 15 |
| Wheat gluten | 5 | 5 | 5 | 5 |
| Corn gluten meal | 5.8 | 5.8 | 5.8 | 5.8 |
| Blood meal, porcine | 5 | 5 | 5 | 5 |
| Corn starch | 3.5 | 0.01 | 2.74 | 2.2 |
| Fish oil, herring | 6.5 | 6.5 | 6.5 | 4.3 |
| Canola oil | 6.6 | 6.6 | 6.6 | 4.4 |
| Vitamin and mineral premix, ADM | 0.5 | 0.5 | 0.5 | 0.5 |
| Vitamin E | 0.5 | 0.5 | 0.5 | 0.5 |
| Choline chloride | 0.3 | 0.3 | 0.3 | 0.3 |
| DL-Met | 0.1 | 0.1 | 0.1 | 0.1 |
| Limestone | 0.1 | 0.1 | 0.1 | 0.1 |
| NaCl | 0.1 | 0.1 | 0.1 | 0.1 |
| CaHPO<sub>4</sub> | 0 | 0.1 | 0.44 | 0 |
| Yttrium oxide | 0.015 | 0.015 | 0.015 | 0.015 |

Ingredients of each diet were pre-weighed in 10kg batches, and all dry ingredients were mixed using a Hobart mixer for 20 min. Fish oil and canola oil were then added to the dry ingredients and mixed for an additional 20 min. Immediately following mixing, diets were pelleted using a laboratory scale steam pellet mill (California Pellet Mill Co., IN, USA) with a 3mm diameter dye to produce pellets of approximately 3 x 5 mm (width x length). Pellets were then oven dried for approximately 16 hours at 40°C using a drying oven. Once dried, the diets were sieved to remove fines and stored at 4°C until the start of the experiment. Samples of each diet were then analyzed for proximate and fatty acid composition (Table 3).

**Table 3.**
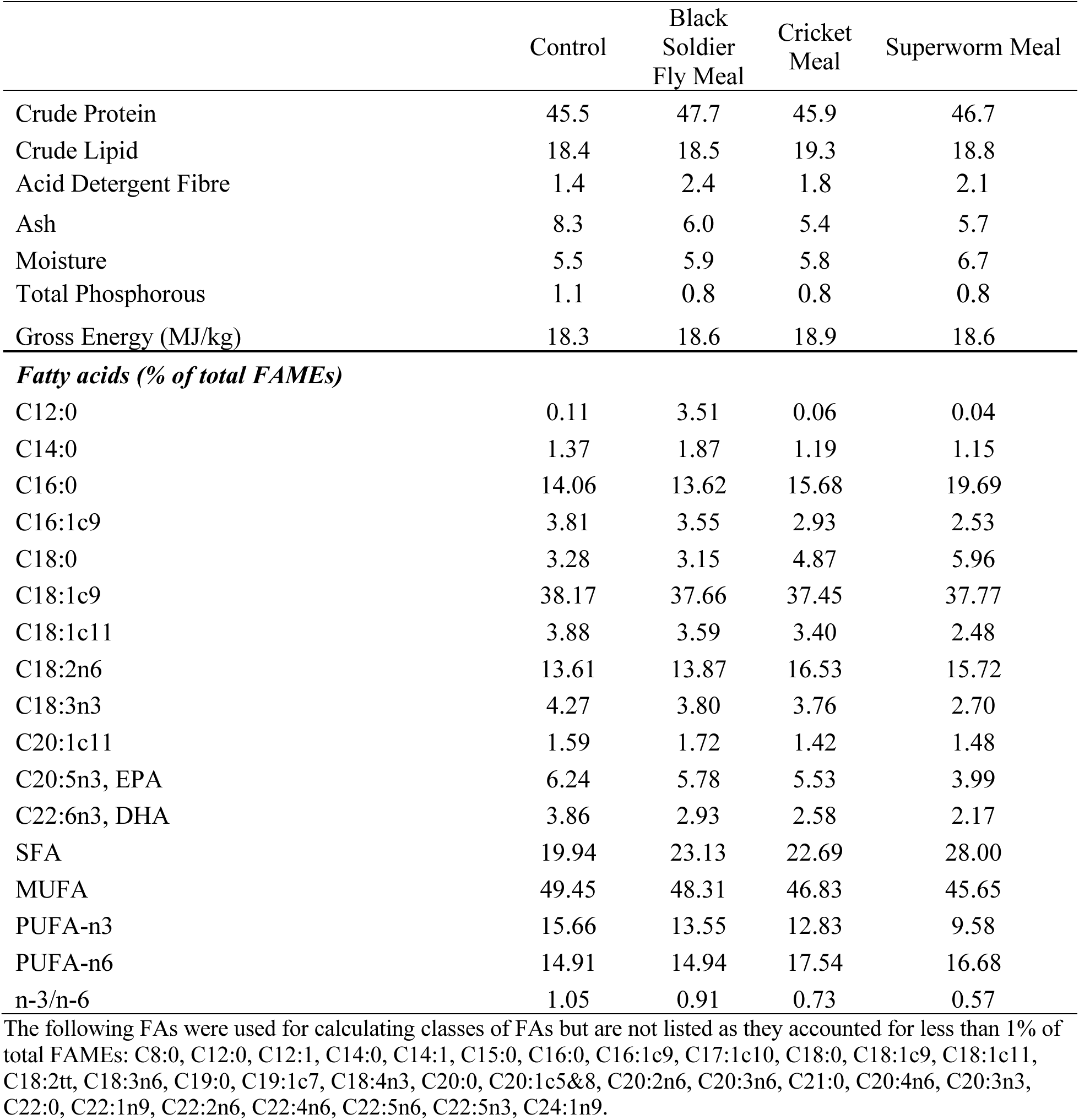
Proximate composition and fatty acid profile reported as % of total Fatty Acid Methyl Esters (FAMEs) of whole experimental diets.

### 2.3 Sample Collection and Calculations

Before trial initiation, 15 fish were collected from the stock population as a pooled sample to determine initial proximate and fatty acid composition of whole-body carcasses. Fish were humanely euthanized using an overdose of tricaine methane sulfonate (MS222; Syndel Canada, Nanaimo, BC, Canada), followed by cervical transection.

During the feeding trial, fish biomass of each tank was bulk weighed every 28 days. At the end of the 84-day growth trial, fish were sedated with MS222 and individual fish weight and fork length of all fish were recorded to determine growth performance. Five fish from each tank were humanely euthanized as above and carcasses were pooled, ground, freeze dried and stored at -20 °C for later proximate and fatty acid analyses (n=3 per treatment). Additionally, viscera and liver of the dissected fish were weighed to determine the viscerosomatic (VSI) and hepatosomatic (HSI) indices (n=3 per treatment). Based on the study by Bureau et al. (2006), the following calculations were used to assess growth performance and nutrient utilization:

- WG (weight gain) = FBW (final body weight) – IBW (initial body weight);
- SGR (specific growth rate, %/day) = ((Ln FBW − Ln IBW) × 100)/ days, where FBW and IBW are the final and initial body weight of fish (g), respectively;
- TGC (thermal growth coefficient) = 100 × (((FBW^1/3^) - (IBW^1/3^))/ (temperature × days));
- Feed conversion ratio (FCR) = feed intake (g)/weight gain (g);
- Condition factor (K) = 100 × (W × FL^3^);
- VSI (viscerosomatic index, %) = (W_V_ × FBW^-1^) × 100;
- HSI (hepatosomatic index, %) = (W_L_ × FBW^-1^) × 100;
- Retention (% of intake): 100 × (FBW × final carcass total nutrient content − IBW × initial total carcass nutrient content)/nutrient intake;
- ADC of nutrient = 100*(1- (% Y in diet/Y in faeces) *(% nutrient in faeces/nutrient in diet)), where Y is yttrium oxide (Y_2_O_3_) added to the diet as a digestibility marker.

To determine nutrient digestibility and faecal nutrient composition, faeces was collected directly from each fish using manual stripping of their abdomen under sedation with MS222 every 28 days. Faecal samples were pooled from each tank of fish (n=3 per treatment) and stored at -20 °C. Faeces from every sampling event was freeze dried and analyzed for phosphorus content. Faeces from the final sampling period (day 84) were analyzed for yttrium, proximate composition, and phosphorous.

### 2.4 Nutritional Analysis

All fish carcasses, faeces, diets, individual fishmeal, black soldier fly meal, cricket meal and superworm meal ingredients were ground and sent to SGS Crop Science Canada for proximate and mineral analyses (Guelph, ON, Canada). Proximate analysis of samples was completed according to AOAC International standards (AOAC, 2003). Dry matter content was determined by drying samples to a constant weight at 135 °C for 2 hr. Crude protein content was determined following the Dumas combustion method with a LECO Analyzer (628 N, LECO Corporation, St. Joseph, MI, USA), using a conversion of N x 6.25. Crude lipid was determined using an ANKOM extractor (ANKOM Technology, Macedon, NY, USA), where the sample was hydrolyzed using HCl solution and extracted using diethyl ether and petroleum ether. Ash was determined by heating samples to 600 °C in a muffle furnace for 2 hr. To determine mineral contents, dry ashed samples were acid digested using HCl, diluted and analyzed by inductively coupled plasma optical emission spectroscopy (ICP-OES; Perkin-Elmer Corporation, Waltham, MA, USA) (Association of Official Analytical Chemists, 2005).

Fatty acid composition of individual fishmeal, insect ingredients, experimental diets, and whole-body carcasses were performed at the University of Guelph department of Human Health and Nutritional Sciences. Total fatty acids were determined by gas liquid chromatography as previously described (Akbari Moghaddam Kakhki et al., 2020) In brief, total lipids were solvent extracted using chloroform and methanol according to the method of Folch (Folch et al., 1957). Then, fatty acids were trans-esterified to fatty acid methyl esters (FAMEs) using boron trifluoride in methanol (14%). FAMEs were analyzed using an Agilent 6890 gas chromatograph (Agilent, Santa Clara, CA, USA) equipped with a flame ionization detector and separated on a fused-silica capillary column (DB-FFAP, cat no. 127-32H2, Agilent, Santa Clara, CA, USA).

### 2.5 Statistical Analysis

All statistical analysis was performed using R statistical software version 4.0.3 (R Core Team, 2023). Results are expressed as means of three replicate tanks and standard deviations. Data for all variables were tested for normal distribution and variance of error using a Shapiro-

Wilk and Levene tests. A one-way analysis of variance (ANOVA) was performed for each response variable. When significant differences were detected in the ANOVA, a post hoc Tukey HSD test was performed to determine where the significant differences were between dietary treatment groups while accounting for multiple testing. P-values less than 0.05 were considered significant.

## 3. Results

### 3.1 Growth Performance

At the end of the 84-day growth trial, the incorporation of black soldier fly, cricket and superworm meal had no significant effect (*p* > 0.05), on final weight, fork length, weight gain, feed intake, condition factor, thermal growth coefficient, specific growth rate, feed conversion ratio, viscerosomatic index, hepatosomatic index, or survival (Table 4).

**Table 4.** Effects of feeding control and insect diets on growth performance, body indices and survival (n=3 per treatment) of rainbow trout after 84 days (mean ± SD).

| Growth Parameter | Control | Black Soldier Fly Meal | Cricket Meal | Superworm Meal | Fishmeal |
| --- | --- | --- | --- | --- | --- |
| Initial weight (g) | 100.8 ± 0.8 | 100.2 ± 0.4 | 99.6 ± 0.3 | 101.3 ± 2.4 | 0.458 |
| Final weight (g) | 290.7 ± 10.1 | 283.9 ± 4.3 | 281.5 ± 9.8 | 282.8 ± 10.8 | 0.628 |
| Fork length (cm) | 27.5 ± 0.3 | 27.4 ± 0.2 | 27.2 ± 0.3 | 27.3 ± 0.2 | 0.454 |
| Weight gain (g/fish) | 189.9 ± 9.8 | 183.8 ± 4.4 | 181.8 ± 9.8 | 181.5 ± 9.9 | 0.642 |
| Feed intake (g/fish) | 174.2 ± 8.6 | 177.6 ± 8.4 | 175.6 ± 9.1 | 172.2 ± 9.3 | 0.894 |
| Condition factor | 1.38 ± 0.01 | 1.37 ± 0.02 | 1.36 ± 0.00 | 1.38 ± 0.02 | 0.208 |
| PER | 2.39 ± 0.01 | 2.17 ± 0.08 | 2.26 ± 0.16 | 2.26 ± 0.07 | 0.111 |
| TGC | 0.28 ± 0.01 | 0.27 ± 0.00 | 0.27 ± 0.01 | 0.27 ± 0.01 | 0.650 |
| SGR | 1.26 ± 0.04 | 1.24 ± 0.01 | 1.24 ± 0.02 | 1.22 ± 0.04 | 0.616 |
| FCR | 0.92 ± 0.01 | 0.97 ± 0.04 | 0.97 ± 0.07 | 0.95 ± 0.03 | 0.449 |
| VSI | 11.5 ± 0.6 | 12.2 ± 0.7 | 12.2 ± 0.8 | 11.5 ± 1.1 | 0.572 |
| HSI | 1.34 ± 0.09 | 1.47 ± 0.05 | 1.39 ± 0.13 | 1.35 ± 0.10 | 0.396 |
| Survival (%) | 100 ± 0.0 | 100 ± 0.0 | 98.7 ± 2.3 | 100 ± 0.0 | 0.441 |

PER; Protein Efficiency Ratio, TGC; thermal growth coefficient, SGR; specific growth rate, FCR; feed conversion ratio, VSI; viscerosomatic index, HSI; hepatosomatic index.

### 3.2 Apparent Digestibility

Statistically significant differences in apparent digestibility coefficients (ADCs) of nutrients and energy were detected among treatments (Table 5). ADC of dry matter was highest in the superworm diet, and lowest in the cricket diet, with control and black soldier fly diets exhibiting intermediate values (p<0.001). ADC of protein was also highest in the superworm diet, but only significantly higher than the black soldier fly and cricket diets (p=0.002). The same trend was also seen in apparent digestibility of energy with superworm being significantly higher than both black soldier fly and cricket diets (p<0.001). The ADC of lipid was not affected by any of the experimental diets (p>0.05). Irrespective of the insect species, ADC of phosphorous was significantly higher in all the insect diets when compared to the control group (p=0.001).

**Table 5.** Effects of feeding control and insect diets on apparent digestibility coefficient (ADC) of key nutrients (n=3 per treatment) for rainbow trout after 84 days (mean ± SD).

| ADC (%) | Control | Black Soldier Fly Meal | Cricket Meal | Superworm Meal | p-value |
| --- | --- | --- | --- | --- | --- |
| Dry Matter | 67.9 ± 0.5<sup>a</sup> | 68.0 ± 0.9<sup>a</sup> | 64.1 ± 0.9<sup>b</sup> | 72.7 ± 1.0<sup>c</sup> | **<0.001** |
| Crude Protein | 81.3 ± 0.4<sup>ab</sup> | 80.6 ± 0.2<sup>a</sup> | 78.1 ± 2.4<sup>a</sup> | 84.5 ± 0.1<sup>b</sup> | **0.002** |
| Crude Lipid | 93.7 ± 5.4 | 95.6 ± 1.3 | 93.0 ± 0.5 | 93.3 ± 0.7 | 0.689 |
| Gross Energy | 76.5 ± 1.3<sup>ab</sup> | 75.6 ± 0.8<sup>b</sup> | 71.4 ± 0.5<sup>c</sup> | 78.2 ± 0.6<sup>a</sup> | **<0.001** |
| Total Phosphorous | 34.7 ± 3.0<sup>a</sup> | 44.7 ± 4.3<sup>b</sup> | 51.2 ± 1.3<sup>b</sup> | 52.7 ± 3.5<sup>b</sup> | **0.001** |

Different superscripts for each response variable indicate a significant difference ( $p < 0.05$ ) between treatments.

### 3.3 Nutrient Retention

Nutrient retention was affected by diet for crude protein and total phosphorous (Table 6). Crude protein retention was highest in the fish meal control group, which was significantly higher than the black soldier fly and cricket treatments (p = 0.001). Crude lipid and gross energy retention were not affected by any of the experimental dietary treatments. Phosphorous retention was improved in the black soldier fly treatment and was significantly higher than the control group (p = 0.023).

**Table 6.** Effects of feeding control and insect diets on nutrient and fatty acid retention (n=3 per treatment) in rainbow trout after 84 days (mean ± SD).

| Retention (% Intake) | Control | Black Soldier Fly Meal | Cricket Meal | Superworm Meal | p-value |
| --- | --- | --- | --- | --- | --- |
| Crude Protein | 40.8 ± 0.5<sup>a</sup> | 34.8 ± 1.1<sup>b</sup> | 37.6 ± 0.8<sup>c</sup> | 38.6 ± 1.0<sup>ac</sup> | **0.001** |
| Crude Lipid | 70.4 ± 4.3 | 77.4 ± 7.2 | 70.8 ± 9.4 | 72.4 ± 4.8 | 0.585 |
| Gross Energy | 43.7 ± 1.8 | 43.9 ± 3.9 | 43.0 ± 3.9 | 43.8 ± 2.0 | 0.977 |
| Total Phosphorous | 39.9 ± 3.0<sup>a</sup> | 54.3 ± 3.7<sup>b</sup> | 51.1 ± 2.8<sup>ab</sup> | 51.5 ± 7.5<sup>ab</sup> | **0.023** |
| ***Fatty acids*** |  |  |  |  |  |
| C12:0, Lauric acid | 14.0 ± 38.9 | 24.2 ± 3.1 | 28.4 ± 54.9 | 82.9 ± 54.5 | 0.279 |
| C14:0, Myristic acid | 28.4 ± 7.8<sup>a</sup> | 42.2 ± 5.2<sup>b</sup> | 36.2 ± 2.1<sup>ab</sup> | 28.4 ± 2.5<sup>a</sup> | **0.024** |
| C16:0 | 31.9 ± 2.9<sup>ab</sup> | 50.2 ± 5.9<sup>c</sup> | 38.3 ± 3.4<sup>a</sup> | 23.6 ± 3.1<sup>b</sup> | **<0.001** |
| C16:1c9, Palmitoleic acid | 32.0 ± 6.6<sup>a</sup> | 55.6 ± 6.0<sup>b</sup> | 44.3 ± 1.7<sup>ab</sup> | 35.8 ± 5.6<sup>a</sup> | **0.003** |
| C18:0 | 36.5 ± 3.7<sup>a</sup> | 58.9 ± 7.6<sup>b</sup> | 38.9 ± 3.0<sup>a</sup> | 22.1 ± 3.9<sup>c</sup> | **<0.001** |
| C18:1c9 | 30.2 ± 4.0<sup>a</sup> | 48.6 ± 5.0<sup>b</sup> | 42.6 ± 4.0<sup>b</sup> | 29.8 ± 1.1<sup>a</sup> | **<0.001** |
| C18:1c11 | 28.0 ± 8.5<sup>a</sup> | 48.0 ± 5.0<sup>b</sup> | 43.0 ± 4.0<sup>ab</sup> | 36.5 ± 7.0<sup>ab</sup> | **0.023** |
| C18:2n6 | 25.7 ± 2.3<sup>ac</sup> | 39.5 ± 4.8<sup>b</sup> | 33.8 ± 2.1<sup>ab</sup> | 21.8 ± 2.5<sup>c</sup> | **<0.001** |
| C18:3n3 | 17.7 ± 5.8<sup>a</sup> | 33.1 ± 4.2<sup>b</sup> | 32.7 ± 4.6<sup>b</sup> | 23.3 ± 5.0<sup>ab</sup> | **0.012** |
| C20:1c11 | 46.7 ± 14.1 | 59.3 ± 8.7 | 58.1 ± 2.8 | 44.2 ± 6.8 | 0.165 |
| C20:5n3, EPA | 11.0 ± 5.1 | 17.6 ± 1.9 | 16.6 ± 1.9 | 12.2 ± 3.5 | 0.114 |
| C22:6n3, DHA | 37.3 ± 14.9<sup>a</sup> | 75.9 ± 11.6<sup>b</sup> | 72.6 ± 6.8<sup>b</sup> | 57.8 ± 9.2<sup>ab</sup> | **0.010** |
| SFA | 31.6 ± 3.0<sup>ac</sup> | 46.0 ± 5.3<sup>b</sup> | 38.8 ± 2.9<sup>ab</sup> | 23.2 ± 2.9<sup>c</sup> | **<0.001** |
| MUFA | 30.4 ± 4.9<sup>a</sup> | 49.1 ± 5.2<sup>b</sup> | 42.9 ± 3.6<sup>b</sup> | 31.0 ± 0.4<sup>a</sup> | **0.001** |
| PUFA-n3 | 21.1 ± 8.3<sup>a</sup> | 37.7 ± 5.1<sup>b</sup> | 35.4 ± 3.1<sup>ab</sup> | 28.0 ± 5.7<sup>ab</sup> | **0.031** |
| PUFA-n6 | 29.0 ± 2.7<sup>ac</sup> | 45.1 ± 5.6<sup>b</sup> | 38.3 ± 2.2<sup>ab</sup> | 24.6 ± 3.0<sup>c</sup> | **<0.001** |

Different superscripts for each response variable indicate a significant difference ( $p < 0.05$ ) between treatments.

Retention of fatty acids in rainbow trout was significantly impacted by the diet (Table 6). As a general trend, retention values for fatty acids were highest in fish fed the black soldier fly diet. Specifically, retention values for C14:0, C16:0, C16:1c9, C18:1c11, C18:2n6, C18:3n3, and C22:6n3 (DHA), were highest in the black soldier fly diet, followed by the cricket diet. Looking at fatty acid classes, the black soldier fly diet displayed a significantly higher retention value for SFA’s compared to the superworm and control diets (p <0.001). MUFA retention was significantly higher in the black soldier fly and cricket diets compared to the superworm and control diets (p = 0.001). PUFA-n3 retention was highest in the black soldier fly group, but only significantly higher than the control (p = 0.031), whereas PUFA-n6 retention was significantly higher in the black soldier fly diet compared to the control and superworm diets (p <0.001).

### 3.4 Whole Body Composition

Final whole-body composition was similar among the dietary treatments, except for crude protein (Table 7). Composition of the carcasses did not differ significantly (p < 0.05) among diets for dry matter, lipid, ash, energy, or phosphorous. Crude protein content was significantly higher in the control and superworm diets (p = 0.022) compared to the black soldier fly and cricket diets. Fatty acid composition of whole-body carcass was altered between dietary treatments, although classes of fatty acids (SFA, MUFA, PUFA n-3 and PUFA n-6) were not significantly different (p < 0.05). Saturated fatty acids, specifically lauric acid and myristic acid, were significantly higher in the whole-body carcass of fish fed the black soldier fly diet compared to all other diets (p < 0.001). Omega-6 PUFA, specifically C18:2n6 (linoleic acid), was significantly higher in the whole-body carcass of fish fed the cricket and superworm diets compared to those fed black soldier fly (p = 0.036). Additionally, MUFA C16:1c9 (palmitoleic acid), was higher in the carcass of fish fed the control and black soldier fly diets compared to the cricket and superworm diets (p = 0.023).

**Table 7.** Effects of feeding control and insect diets on whole-body composition and fatty acid profile (n=3 per treatment) of rainbow trout after 84 days (mean ± SD).

| Whole-Body Composition | Control | Black Soldier Fly Meal | Cricket Meal | Superworm Meal | p-value |
| --- | --- | --- | --- | --- | --- |
| Dry Matter (% of carcass) | 30.3 ± 0.4 | 31.3 ± 0.3 | 31.1 ± 0.5 | 31.1 ± 0.4 | 0.092 |
| Crude Protein (% of carcass) | 16.70 ± 0.13<sup>a</sup> | 16.02 ± 0.09<sup>b</sup> | 16.43 ± 0.43<sup>ab</sup> | 16.71 ± 0.08<sup>a</sup> | **0.022** |
| Crude Lipid (% of carcass) | 11.37 ± 0.50 | 12.62 ± 0.47 | 12.18 ± 0.59 | 12.01 ± 0.49 | 0.094 |
| Ash (% of carcass) | 2.88 ± 0.09 | 2.85 ± 0.11 | 2.91 ± 0.08 | 2.80 ± 0.07 | 0.471 |
| Gross Energy (MJ/kg of carcass) | 7.1 ± 0.20 | 7.4 ± 0.18 | 7.4 ± 0.23 | 7.3 ± 0.18 | 0.197 |
| Phosphorous (% of carcass) | 0.4 ± 0.02 | 0.4 ± 0.03 | 0.4 ± 0.03 | 0.3 ± 0.04 | 0.649 |
| ***Fatty acids (% of total FAMES)*** |  |  |  |  |  |
| C12:0 | 0.06 ± 0.10<sup>a</sup> | 1.31 ± 0.05<sup>b</sup> | 0.05 ± 0.05<sup>a</sup> | 0.11 ± 0.06<sup>a</sup> | **<0.001** |
| C14:0 | 1.64 ± 0.10<sup>a</sup> | 1.88 ± 0.01<sup>b</sup> | 1.49 ± 0.02<sup>a</sup> | 1.56 ± 0.10<sup>a</sup> | **<0.001** |
| C16:0 | 15.85 ± 1.23 | 15.27 ± 0.20 | 15.42 ± 0.27 | 16.76 ± 1.24 | 0.242 |
| C16:1c9 | 4.91 ± 0.08<sup>a</sup> | 4.91 ± 0.09<sup>a</sup> | 4.32 ± 0.22<sup>b</sup> | 4.34 ± 0.42<sup>b</sup> | **0.023** |
| C18:0 | 4.30 ± 0.53 | 4.18 ± 0.04 | 4.66 ± 0.08 | 4.72 ± 0.50 | 0.275 |
| C18:1c9 | 36.32 ± 0.77 | 36.50 ± 0.30 | 36.08 ± 0.83 | 37.06 ± 0.61 | 0.373 |
| C18:1c11 | 3.43 ± 0.44 | 3.48 ± 0.03 | 3.38 ± 0.07 | 3.17 ± 0.47 | 0.669 |
| C18:2n6 | 14.17 ± 0.81<sup>ab</sup> | 13.80 ± 0.14<sup>a</sup> | 15.46 ± 0.22<sup>b</sup> | 14.45 ± 0.75<sup>b</sup> | **0.036** |
| C18:3n3 | 2.24 ± 0.38 | 2.38 ± 0.06 | 2.55 ± 0.23 | 2.05 ± 0.36 | 0.251 |
| C20:1c11 | 2.48 ± 0.28 | 2.26 ± 0.05 | 2.17 ± 0.04 | 2.39 ± 0.28 | 0.308 |
| C20:5n3, EPA | 2.37 ± 0.54 | 2.29 ± 0.08 | 2.36 ± 0.09 | 2.02 ± 0.37 | 0.569 |
| C22:6n3, DHA | 5.11 ± 0.91 | 5.04 ± 0.26 | 4.98 ± 0.10 | 4.92 ± 0.57 | 0.979 |
| SFA | 22.50 ± 1.5 | 23.19 ± 0.18 | 22.53 ± 0.42 | 23.57 ± 1.6 | 0.602 |
| MUFA | 43.73 ± 0.35 | 43.66 ± 0.22 | 42.92 ± 0.22 | 44.10 ± 0.75 | 0.207 |
| PUFA-n3 | 11.20 ± 2.07 | 11.09 ± 0.42 | 11.23 ± 0.11 | 10.27 ± 1.51 | 0.768 |
| PUFA-n6 | 17.57 ± 0.90 | 17.06 ± 0.04 | 18.85 ± 0.33 | 17.63 ± 0.96 | 0.059 |
| n-3/n-6 | 0.64 ± 0.14 | 0.65 ± 0.02 | 0.60 ± 0.01 | 0.59 ± 0.12 | 0.807 |

<sup>1</sup> Initial body composition: dry matter 20.7 %, protein 16.0%, lipid 10.4%, ash 0.4%, phosphorous 0.3% and energy 6.6 MJ/kg. Different superscripts for each response variable indicate a significant difference (p<0.05) between treatments. The following FAs were used for calculating classes of FAs but are not listed as they accounted for less than 1% of total FAMES: C14:1, C15:0, C18:2t, C18:3n6, C19:0, C18:4n3, C20:0, C20:1c5&8, C20:2n6, C20:3n6, C21:0, C20:4n6, C20:3n3, C22:0, C22:1n9, C22:2n6, C22:4n6, C22:5n6, C22:5n3, C24:0, C24:1n9.

## 4. Discussion

Incorporating new ingredients, such as insect meals, into the commercial diets of farmed fish requires comprehensive evaluation on their possible effects on fish growth performance, nutritional composition, and environmental impacts. Currently, there has been a significant amount of research carried out on the effect of black soldier fly meal in salmonid diets, yet there is little comparison to alternative sources of insect protein. To the best of the authors knowledge, this was the first study to compare black soldier fly, cricket and superworm meals at the 15% dietary inclusion level in farmed rainbow trout. Encouragingly, no adverse effects on growth performance or survival were observed at this level of inclusion for any of the experimental diets (Table 4). These findings align with previous studies that have demonstrated the capacity to replace fishmeal with black soldier fly at inclusion levels upwards of 15% without negatively impacting fish growth performance (Bordignon et al., 2022; Józefiak et al., 2019; Terova et al., 2019). There were, however, differences among treatments in other factors measured in this study.

It is important to note that the impact of different insect meals on rainbow trout growth varies widely among studies. For instance, Józefiak et al. (2019) reported that a 20% inclusion of cricket (*Gryllodes sigillatu*s) had negative effects on the final average body weight of rainbow trout. In contrast, another study found no negative impact on growth performance when 25% of the diets’ gross energy was replaced with live adult house cricket (*Acheta domestica*) and superworm (Turek et al., 2020). At 11% inclusion, superworm showed no adverse effects on rainbow trout growth, but growth rates were reduced at a 22% inclusion rate (Shekarabi et al., 2021). These observations suggest that the appropriate dietary inclusion rate for insect meals could be contingent on the specific insect species, the overall diet formulation, and the conditions under which the studies were executed. The advantage of the present study is that it allows for direct comparison among treatments, as treatments were run in parallel under the same conditions. Considering the results of this study alongside previous feeding trials involving black soldier fly, cricket and superworm, a 15% inclusion rate for all insects used appears to fall within an acceptable dietary range for rainbow trout.

However, further research is needed to optimize the use of insect meals in rainbow trout diets, including exploring different inclusion levels and combinations of insect species, to fully harness their potential as sustainable and effective feed alternatives for the aquaculture industry.

The insect treatments had significant impacts on the digestibility of dry matter, crude protein, gross energy and total phosphorous (Table 5). Previous studies have found that high inclusion of insect protein in the diet of rainbow trout has a significant effect on nutrient digestibility. In a study evaluating the incorporation of yellow mealworm *(Tenebrio molitor*) in the diet of rainbow trout it was noted that 50% dietary inclusion of insect meal resulted in decreased protein digestibility compared to a 25% inclusion (Belforti et al., 2015).

Comparatively, another study examining the effects of feeding yellow mealworm to rainbow trout found that up to 25% total inclusion of the insect protein had no effects on nutrient digestibility (Rema et al., 2019). A study that fed crickets to olive flounder (*Paralichthys olivaceus*) found reduced protein digestibility when more than 20% of fishmeal was replaced (Jeong et al., 2021). In the current study, fish fed the cricket diet had significantly lower protein and energy digestibility compared to the control and other insect diets (Table 5), which may reduce its potential as an alternative to fishmeal.

The black soldier fly and cricket diets showed lower crude protein digestibility compared to the superworm and control diets (Table 5), which may indicate superworms have a higher potential as a fishmeal and fish oil replacement using this metric as an indicator. Previous research into cricket meal inclusion in largemouth bass (*Micropterus salmoides*) diets has found that 36% total dietary inclusion and 60% replacement of fishmeal with cricket meal resulted in lowered crude protein content in the whole-body of fish (Wang et al., 2022).

However, in the current study the superworm meal ingredient had the lowest content of protein (46 vs 55-66%) and thus replaced far less fishmeal (47 vs 58-74% replacement) compared to the black soldier fly and cricket diets (Table 1), which may explain this result. Like black soldier fly larvae, superworm could be defatted to increase its protein content and reduce its lipid content, which was much higher than the two other insect meal ingredients (40 vs 11-19% lipid). In contrast, the superworm diet replaced more fish oil and the lack of negative effects on growth performance indicate the remaining fish oil in the diet met the requirements for essential omega-3 PUFA. This result indicates that superworm can replace a combination of fishmeal and fish oil without negative effects on growth performance, while maintaining protein digestibility where black soldier fly and cricket diets did not.

Subtle differences in nutrient retention were detected in this study, where crude protein retention was reduced by the dietary inclusion of black soldier fly and cricket diets (Table 6). This may be linked to reduce protein and amino acid digestibility. Similar results were found in a previous study where rainbow trout were fed diets containing 28% defatted black soldier fly and experienced decreased protein utilization efficiency (Stadtlander et al., 2017). These results, however, are in contrast to a study where it was found that up to 25% protein substitution with black soldier fly meal in the diets of rainbow trout and Atlantic salmon did not significantly impact protein retention (Weththasinghe et al., 2021). Although reduced protein retention did not negatively impact fish growth in our 12-week trial, there is potential that decreased protein retention in a longer commercial production cycle could reduce overall growth performance.

There was high variability among the fatty acid profiles of the individual insect meals and fishmeal (Table 1) yet the fatty acid composition of the whole-body of fish fed the insect diets were not significantly altered in terms of total saturated fatty acids (SFA), monounsaturated fatty acids (MUFA), omega-3 and omega-6 polyunsaturated fatty acids (PUFA; Table 7). The experimental diets were specifically formulated to meet the omega-3 PUFA requirements of rainbow trout, and as such at least 4.3% of the diet was fish oil (Table 2). While insect ingredients were higher in SFA’s in comparison to fishmeal (43-59% vs 28% total FAMEs; Table 1), the adjustment of fishmeal and fish oil in the diet seemed to compensate for this difference and the resulting total SFA’s in the whole carcasses of the experimental fish were all within a 1.5% difference. While total SFA’s in the whole carcass of the fish were not significantly impacted, it was found that SFA C12:0, lauric acid, was significantly higher in the black soldier fly treatment group (p < 0.001). Similar results have been found in previous studies where inclusion of black soldier fly larvae in the diet of rainbow trout has significantly increased lauric acid in the fillet (Bruni et al., 2020).

Interestingly, both the ingredients and whole-body carcasses of the rainbow trout control, cricket and superworm diets had similar content of lauric acid and SFA’s in (Table1, Table 7). Additionally, cricket superworm meals had a high content of linoleic acid (Table 1), which was directly correlated to a higher carcass content in fish fed the cricket and superworm diets when compared to the black soldier fly treatment (Table 7). Linoleic acid is an essential fatty acid for humans which cannot be synthesized by the body, and dietary supplementation has been related to improved blood cholesterol levels (Dhull et al., 2021). Crickets and mealworm have previously been reported as a good source of linoleic acid (Alfiko et al., 2022; Pilco-Romero et al., 2023), and the current study found a direct reflection of this fatty acid increased in the carcass of fish fed cricket. Compared to black soldier fly, reduced content of SFA’s in crickets and superworms, higher linoleic acid contents, as well as lack of significant effects on the fatty acid profile of fish demonstrates crickets and superworms have higher potential as fishmeal alternatives in diets for rainbow trout when considering these factors.

Omega-3 PUFA’s were also significantly higher in the fishmeal ingredient compared to all three insect meal ingredients (30% vs 0.9-2% total FAMEs; Table 1), yet this did not significantly impact whole carcass omega-3 content, likely due to the level of fishmeal replacement in the diet. This may in part be because fish are able to synthesize EPA (eicosapentaenoic acid 20:5n-3) and DHA (docosahexaenoic acid 22:6n-3) from ALA (Alpha-Linoleic Acid, 18:3n3), which was present in all the experimental diets (Alfiko et al., 2022). Studies that have examined the effects of insect meal on omega-3 PUFA’s in muscle tissue of rainbow trout have garnered mixed results. Some studies have found that insect meal inclusion in the diets of rainbow trout results in decreased omega-3 PUFA’s in fish muscle tissue (Belforti et al., 2015; Fabrikov et al., 2021; Mancini et al., 2018; Turek et al., 2020). However, in agreement with the results of our study, Bruni et al. (2020) included 10.5 and 21% full fat black soldier fly meal with respective substitution of 25 and 50% fishmeal and found no significant impacts on the fillet omega-3 PUFA composition (Bruni et al., 2020). While there may be subtle differences among our treatments with respect to fatty acid profiles, our results indicate that fatty acid composition is quite comparable among treatments.

To ensure the long-term viability of the aquaculture industry, improving nutrient utilization by fish is crucial. In freshwater ecosystems, phosphorus loading is linked to elevated primary productivity and eutrophication (Bureau & Hua, 2010; Schindler, 1977). Researchers have highlighted the potential of optimizing digestible phosphorus in aquaculture diets to reduce phosphorus waste and the potential for harmful algal blooms (Cho & Bureau, 2001). In the current study, diets were formulated to have the same dietary phosphorus content, but final feed analysis revealed that the control had 1.1% total phosphorus, while the insect diets contained 0.8% total phosphorus. Insect meals were lower in phosphorus compared to fishmeal ingredients, which may make them a suitable dietary protein and lipid source for aquafeeds aiming to optimize phosphorus utilization. Excess dietary phosphorus is excreted as soluble and solid waste (Dauda et al., 2019). Our results suggest that incorporating 15% superworm meal, adult cricket meal, and black soldier fly meal significantly improved apparent phosphorus digestibility (Table 5). Additionally, insect inclusion resulted in higher phosphorus retention, with black soldier fly meal showing significantly higher retention than the control. While eliminating nutrient loading from aquaculture is unattainable, strategies such as using highly digestible and low-phosphorus ingredients in fish feeds can mitigate excessive phosphorus excretion (Coloso et al., 2003). Our findings align with previous studies that found silkworm (*Bombyx mori L*.)-based diets enhanced phosphorus digestibility in diets for Pacific white shrimp (*Litopenaeus vannamei*), and housefly (*Musca domestica*) maggot meal reduced phosphate levels in the tank water of Nile tilapia (*Oreochromis niloticus*) (Rahimnejad et al., 2019; Wang et al., 2017). The aquaculture industry would benefit from further research on effects of insect species at various inclusion levels in relation to nutrient utilization to enhance our understanding of the potential for insect meal to replace fishmeal and fish oil while reducing environmental impacts from rainbow trout farming.

## 5. Conclusions

This study revealed that 15% dietary inclusion of black soldier flies, adult crickets, and superworms were effective in partially replacing fishmeal and fish oil in diets for rainbow trout. In comparison to the control diet, all three insect diets performed similarly in terms of fish growth, body indices and survival. Compared to the control, black soldier flies and crickets reduced protein content and retention in the whole-body, which was not found for superworms. There was no effect of insects on total SFA, MUFA, PUFA n-3 and PUFA n-6, in the whole-body, yet there was an increase in lauric acid and myristic acid when fed black soldier flies and increased linoleic acid when fed cricket and superworm compared to black soldier fly. Additionally, all three insect diets improved apparent digestibility of phosphorous. These results provide further support for black soldier fly, adult cricket and superworm as sustainable ingredients in rainbow trout diets with potential to reduce phosphorus loading on the environment as well.

## Author Contributions

Sonja Drosdowech conducted the feeding trial, data analysis and wrote the original draft of the manuscript. Neil Rooney acquired funding, contributed to project administration, review and editing. Marcia Chiasson, David Huyben and David Ma contributed to project planning, review and editing.

## Acknowledgements

Funding for the use of facilities at the Ontario Aquaculture Research Centre was provided by the Ontario Ministry of Agriculture, Food and Rural Affairs (OMAFRA). The authors thank Fisheries and Oceans Canada (DFO) as well as Entomo Farms for their ingredient and financial contributions to this research project. The preliminary findings of this study were previously presented at the annual conference for the International Association of Great Lakes Research. The authors thank the staff at the OARC, Michael Kirk, Anne Easton, Graeme Woodall and Wesley Chase, as well as student volunteers, Reilly O’Connor, Maya Persram, Zachary Jones, Carmi Riesenbach, Junyu Zhang, Laina Weiss, and Lyle Novella-Talusan for help with sampling

